# Integration of polarization and intensity contrast information in a highly visual animal

**DOI:** 10.64898/2026.08.09.743780

**Authors:** Verónica Pérez-Schuster, Lucca Salomon, Tomás Manuel Chialina, Juliana Reves Szemere, Federico Sevlever, Gabriela Hermitte, Martín Berón de Astrada

## Abstract

Polarization vision subserves diverse biological functions, such as navigation, communication and target-motion detection. Regarding the detection of biological targets, studies on semi-terrestrial crabs suggest that polarization and intensity contrast are processed in separate visual channels. Information about the polarization contrast of targets would be extracted independently of intensity contrast, and the two signals combined downstream in the visual system. However, understanding how the information about these visual attributes is processed and integrated has been limited, as it is technically challenging to present visual stimuli in which both the polarization and the intensity contrast of a stimulus are controlled. Here we developed a monitor screen that allows us to present stimuli in which both contrasts can be controlled. Thus, to study how polarization and intensity information is processed to increase target detection, we presented moving stimuli with controlled polarization and intensity contrast while recording the cardiac response of the semi-terrestrial crab *Neohelice granulata* as a sensitive readout of its visual perception. Our results suggest that *Neohelice* possesses similar sensitivity to vertically and horizontally polarized light; thus, previously reported responses of the animals to polarized stimuli in which figure and background have the same intensity are likely accounted for by the comparison of two polarization channels. In addition, we determined that a moving polarization-only stimulus has a salience equivalent to that of an intensity-only stimulus with a Michelson contrast of 0.51. Finally, we studied how polarization and intensity contrast information is integrated, and found that polarization contrast increases the salience of an intensity-contrast-based target mostly when its intensity contrast is low, i.e. when information about intensity contrast is more ambiguous.

## INTRODUCTION

Unlike vertebrates, many invertebrates make use of the linear polarization of light as a source of valuable visual information. Arthoropods and Cephalopods use polarization information with highly different purposes. Polarization vision provides insects with a visual compass (R. Wehner and Müller (2006)), allows flying insects to detect water bodies (Wildermuth (1998)), serves as a communication channel in cephalopods and crustaceans (Gagnon et al. (2015)), and is used as a source of information to enhance visual contrast in cephalopods and different arthropod species (Shashar, Rutledge, and T. Cronin (1996), Cartron et al. (2013), Sharkey, Partridge, and N. Roberts (2015), and Smithers (2019)).

Polarization sensitivity in invertebrate photoreceptors derives from the differential absorption of polarized light in the microvilli that form the light-sensitive structure of the cell. Because rhodopsin is a dichroic photopigment and its distribution in the microvilli is on average spatially constrained, light absorption is maximal when the e-vector of the incident light is parallel to the long axis of the microvilli (reviewed in N. W. Roberts, Porter, and T. W. Cronin (2011)). The simplest photoreceptor arrangement that serves to analyze polarization in a given viewing direction involves a pair of photoreceptors with orthogonal e-vector tuning axes (Labhart (2016)). No polarization vision system based solely on two orthogonal analyzers can provide sufficient information to determine the angle and the degree of polarization of the incident light. However, any such system can still provide strong polarization contrast information between a figure and a background with different polarizations, except for certain combinations of figure and background intensities and polarizations (M. J. How and N. J. Marshall (2014)).

Cephalopods and decapod crustaceans have been shown to possess exceptional polarization vision. In both groups of animals, orthogonal polarization analyzers are present across the entire retina; thus, polarization contrast can be analyzed throughout the animal’s entire visual field. (R. Glantz (2007), Talbot and J. N. Marshall (2011), and Alkaladi, M. How, and J. Zeil (2013)). In cephalopods, it was shown that polarization vision can be used for object discrimination, the detection of object motion, to improve vision in turbid waters and as a communication channel (Shashar, Rutledge, and T. Cronin (1996), Temple et al. (2012), and Tuthill and Johnsen (2006)). In crayfish, polarization vision has been shown to mediate optomotor and evasive responses (R. M. Glantz and Schroeter (2007) and Tuthill and Johnsen (2006)). In crabs, polarization vision has mainly been studied in the context of target detection, where it was found to be involved in detecting the motion of visual danger stimuli, and it has been suggested to mediate the detection of conspecifics. (M. How, Pignatelli, et al. (2012), M. How, Christy, et al. (2015), and Jochen Zeil and Hofmann (2001)).

Although there is compelling evidence that different animal species use polarization information to enhance object and motion vision, how animals combine polarization information with intensity information is an open question (discussed in Labhart (2016)). One explanation is that, for a viewing direction, polarization information is initially combined with luminance information and then compared with an equivalent measure for another viewing direction to extract a single measure of visual contrast. Another explanation is that polarization contrast is extracted in a channel separate from that of intensity contrast, and the two contrast signals are then combined downstream in the nervous system (Smithers (2019) and Smithers, Brett, and M. How (2024)). The theoretical framework proposed for the processing of polarization contrast independently of the intensity contrast channel is based on color vision models (Bernard and Rüdiger Wehner (1977) and M. J. How and N. J. Marshall (2014)). Briefly, these models propose that, in a two-channel polarization vision system, the inputs from the two types of orthogonal photoreceptors are combined in an opponent manner in a first-order neuron. Subsequently, the activity of two of these neurons, one viewing the object and one the background, would be compared in a second-order neuron, the activity of which would code for the information about polarization contrast. Laboratory and field studies performed in different semiterrestrial crabs have provided empirical support for such a model of polarization contrast perception (e.g. Basnak et al. (2018), M. How, Christy, et al. (2015), Smithers (2019), and J. Zeil (2023)).

Here, we aimed to study how polarization and intensity contrast are integrated in a highly polarization-sensitive crab,*Neohelice granulata* (Basnak et al. (2018)). *Neohelice* inhabits shallow intertidal and semiterrestrial environments. In these habitats, light arising from the sky and from reflections off water surfaces and wet sand is highly polarized (J. Zeil (2023)). Polarization vision has been shown to increase semi-terrestrial crabs’ ability to detect moving targets in their habitat (M. How, Christy, et al. (2015)). Under laboratory conditions, by studying the escape responses of *Neohelice* to visual danger stimuli of different polarization contrasts, we found that the intensities of the escape responses were consistent with a polarization contrast detection system based on two channels: one sensitive to vertically polarized light and the other to horizontally polarized light (Basnak et al. (2018)). However, the study of the integration of polarization and intensity information in our and other animal models has been limited by methodological issues (but see Smithers (2019)). Most of the recent studies on polarization vision have taken advantage of the possibility to present highly salient polarized stimuli in modified LCD screens (e.g. Temple et al. (2012), M. How, Pignatelli, et al. (2012), and Basnak et al. (2018)). These screens permit control of the angle and degree of polarization of light while keeping the intensity and spectral content of a virtual figure and its background equal. However, because these screens do not allow the intensity of the emitted light to be controlled, the intensity contrast of the cast images cannot be tuned. Here we modified a 3D monitor screen to cast vertically and/or horizontally polarized light, i.e., aligned with the two orthogonal polarization channels of *Neohelice*, while also allowing intensity contrast to be controlled. Thus, we presented stimuli with controlled polarization and intensity contrast and recorded the animals’ cardiac response as a sensitive readout of their visual perception (Pérez-Schuster et al. 2023). Our results indicate that the proposed vertical and horizontal polarization channels of *Neohelice* have similar light sensitivity; thus, the responses previously observed to polarized stimuli in which figure and background have equal intensity cannot be explained by differential perception of vertically and horizontally polarized light. In addition, we determined that a moving polarization-only stimulus has a salience equivalent to that of an intensity-only stimulus with a Michelson contrast of 0.51. Finally, we studied how polarization and intensity contrast integrate to give rise to the perceived contrast, and found that polarization contrast increases the salience of an intensity-contrast stimulus particularly when its intensity contrast is low.

## RESULTS

Most recent studies on object-based polarization vision in Crustacea have been conducted by presenting a moving figure of equal intensity and spectral composition to the background, but differing in its polarization content (e.g. Temple et al. (2012), M. How, Pignatelli, et al. (2012), and Basnak et al. (2018)). By using such methodology, we and other authors concluded that different crab species possess two polarization channels aligned with the vertical and horizontal orientations (M. How, Pignatelli, et al. (2012) and Basnak et al. (2018)). However, morphological studies performed in fiddler crabs have shown that, at the eye equator, the area of the eye where we have studied polarization sensitivity in *Neohelice*, the rhabdoms are enriched with vertically oriented microvilli relative to horizontally oriented ones (Alkaladi, M. How, and J. Zeil (2013)). Thus, the ommatidia in this region of the eye could be more sensitive to vertically than to horizontally polarized light. If this were the case, the detection of a vertically or horizontally polarized figure over a background with different polarization but equal intensity could simply be the consequence of differential light absorption by the retinula cells, which would give rise to intensity contrast perception. Here we developed a monitor screen that allows controlling both the intensity of the emitted light and the amount of vertically/horizontally linearly polarized light (i.e. the orientations matching the orientation of the two polarization channels of *Neohelice* (Basnak et al. (2018))(see Intensity and polarization screen). Taking advantage of the stimulation possibilities offered by this monitor, we performed two experiments to explore whether the responsiveness to polarization contrast observed in *Neohelice* is the consequence of differential sensitivity to vertically and horizontally polarized light. The first experiment aimed to investigate whether *Neohelice* exhibits differential sensitivity to light with different polarization orientations. If animals respond differentially to a dark target moving over dissimilar polarized backgrounds of equal intensity, this would suggest that they perceive the backgrounds differently. To this purpose we presented different stimuli to tethered crabs and recorded their cardiac activity as an indicator of their sensory perception (Fig. 2;Pérez-Schuster et al. (2023)). In the three main experimental groups of this experiment (shown in pink shades in Fig. 2, a dark edge advanced over a vertically polarized background, a horizontally polarized background, or a non-polarized background (equal content of vertically and horizontally polarized light). All the backgrounds presented the same light intensity. table S1 summarizes the main characteristics of the visual stimuli, the intensity contrast and the polarization contrast of the stimuli. The temporal profile of the cardiac activity along the stimulation trials is shown in Fig. 2A. The light blue area highlights the moment of visual stimulation. The quantification of the cardiac responses the area under the curve during the stimulation period for the different groups is shown in Fig. 2B. Independently of the background condition, we found no difference in the cardiac response to the dark edge, suggesting that the different backgrounds were perceived similarly (pinkish groups: Vertical_B_ = 0.8*±*0.1, Horizontal_B_ = 0.8*±*0.1; Non-POL_B_ = 0.9*±*0.1; Tukey tests: Horizontal_B_ vs Non-POL_B_ p=0.21, Horizontal_B_ vs Vertical_B_ p=0.71, Vertical_B_ vs Non-POL_B_ p=0.81; N = 48). In addition, the experiment included a further group run throughout the present study: a fully vertically polarized edge moving over a fully horizontally polarized background of equal intensity (green curve in Fig. 2; cardiac response Vertical_E_-Horizontal_B_ = 0.53*±*0.05). The response to this reference stimulus indicate that the temporal profile and the magnitude of the cardiac response to polarized stimuli are robust across experiments (see Figs. 2 to 4).

**Figure 1.**
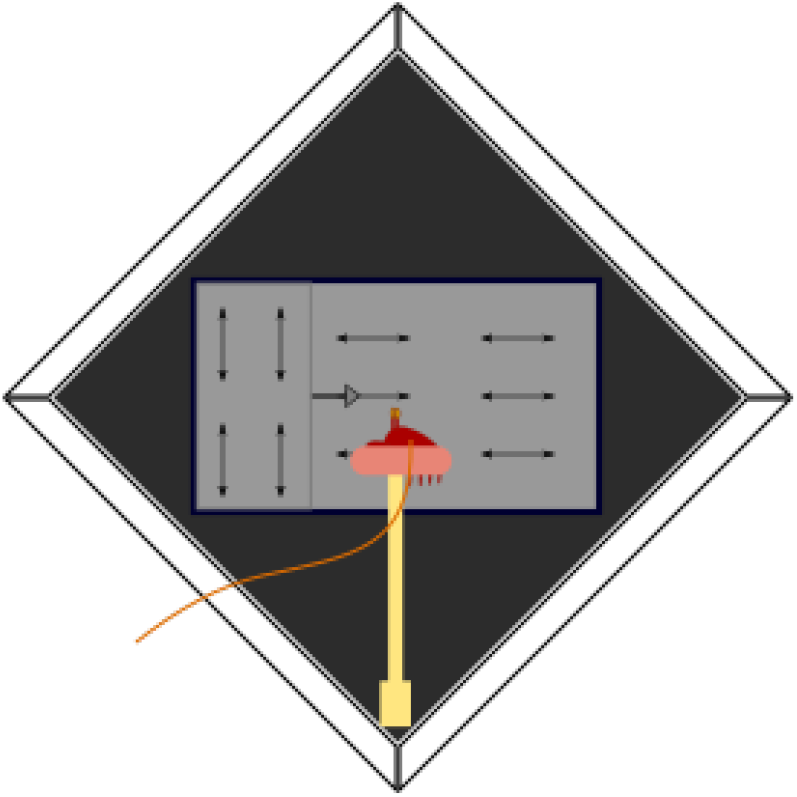
Experimental setup. The crabs were held in place with an adjustable clamp. Their claws and legs were secured against the body with a rubber band. Crabs were positioned 20 cm away from the monitor screen, with the lateral pole of the eye looking at its centre. Vertical edge stimuli translating horizontally across the monitor screen were presented to the animals.

**Figure 2.**
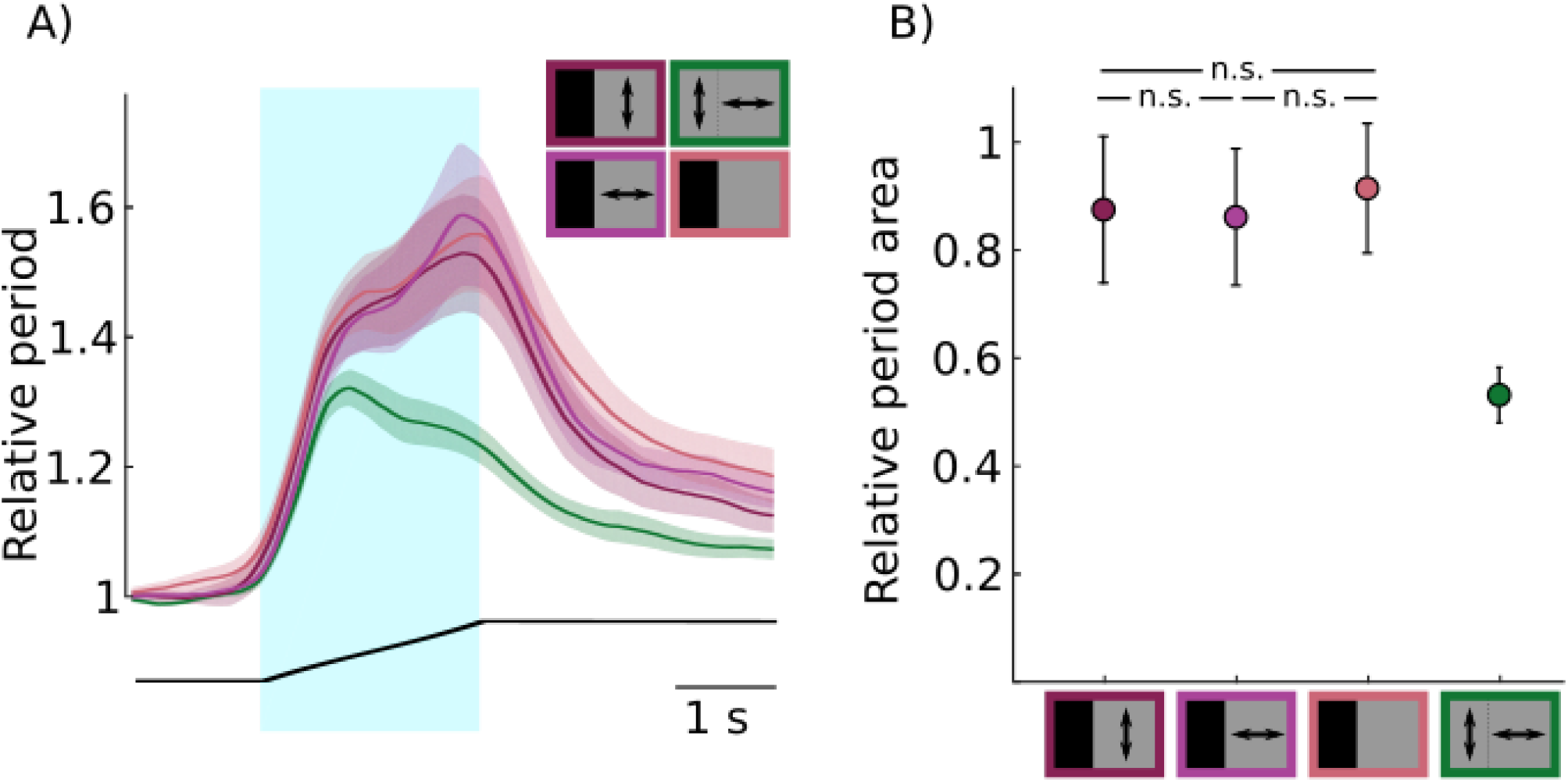
Cardiac responses of *Neohelice* to different background polarizations. (A) Animals’ relative cardiac period (mean±s.e.m., N = 48) as a function of time in response to a moving edge. The light blue shaded area indicates the time of presentation of the stimulus. The black line below the plots illustrates the horizontal position of the moving edge in the screen. The stimulus consisted of a dark edge translating over a vertically polarized, horizontally polarized, or non-polarized background (pinkish curves), and a vertically polarized edge moving over a horizontally polarized background of the same light intensity (green curve). (B) Relative period area integrated over the stimulation time (mean±s.e.m., N = 48). Specific characteristics of the stimuli are detailed in table S1. Statistical differences are indicated as follows: n.s., non-significant differences.

To further study if the response to polarized contrast stimuli are the consequence of differential sensitivity to vertically and horizontally polarized light we performed the experiment shown in Fig. 3. We presented two stimuli with polarization contrast but no intensity contrast: a vertically polarized edge moving over a horizontally polarized background and a horizontally polarized edge moving over a vertically polarized background (shown in green shades in the figure; see table S2 for more stimuli details). We knew from previous studies that crabs’ escape response to intensity contrast stimuli of opposite polarity are highly asymmetrical. Dark edges moving over light backgrounds evoke robust escape responses, whereas light edges moving over dark backgrounds evoke virtually no escape response (Bengochea (2017)). Thus, if the polarization contrast sensitivity observed when presenting edges and backgrounds of the same intensity is simply the consequence of differential sensitivity to vertically and horizontally polarized light, it is expected that such stimuli with opposite polarization contrast evoke asymmetrical responses. However, cardiac responses to stimuli of equal edgebackground intensity but opposite polarization contrast were highly similar (cardiac response of VerticalE-HorizontalB = 0.7*±*0.1 and of Horizontal_E_-Vertical_B_ = 0.5*±*0.1; Tukey test: Vertical_E_-Horizontal_B_ vs Horizontal_E_-Vertical_B_ p=0.06; N = 24). The response of both groups resulted significantly higher than a control group in which a horizontally polarized edge translates over a background of same intensity and polarization (yellow curve in Fig. 3A; cardiac response of Horizontal_E_-Horizontal_B_ = 0.12*±*0.03; statistics not shown). The results of the previous experiments indicate that *Neohelice* possesses similar sensitivity to horizontally and vertically polarized light; thus, the responses to stimuli in which figure and background have equal intensity but differ in polarization do not appear to be explained in terms of intensity contrast perception.

**Figure 3.**
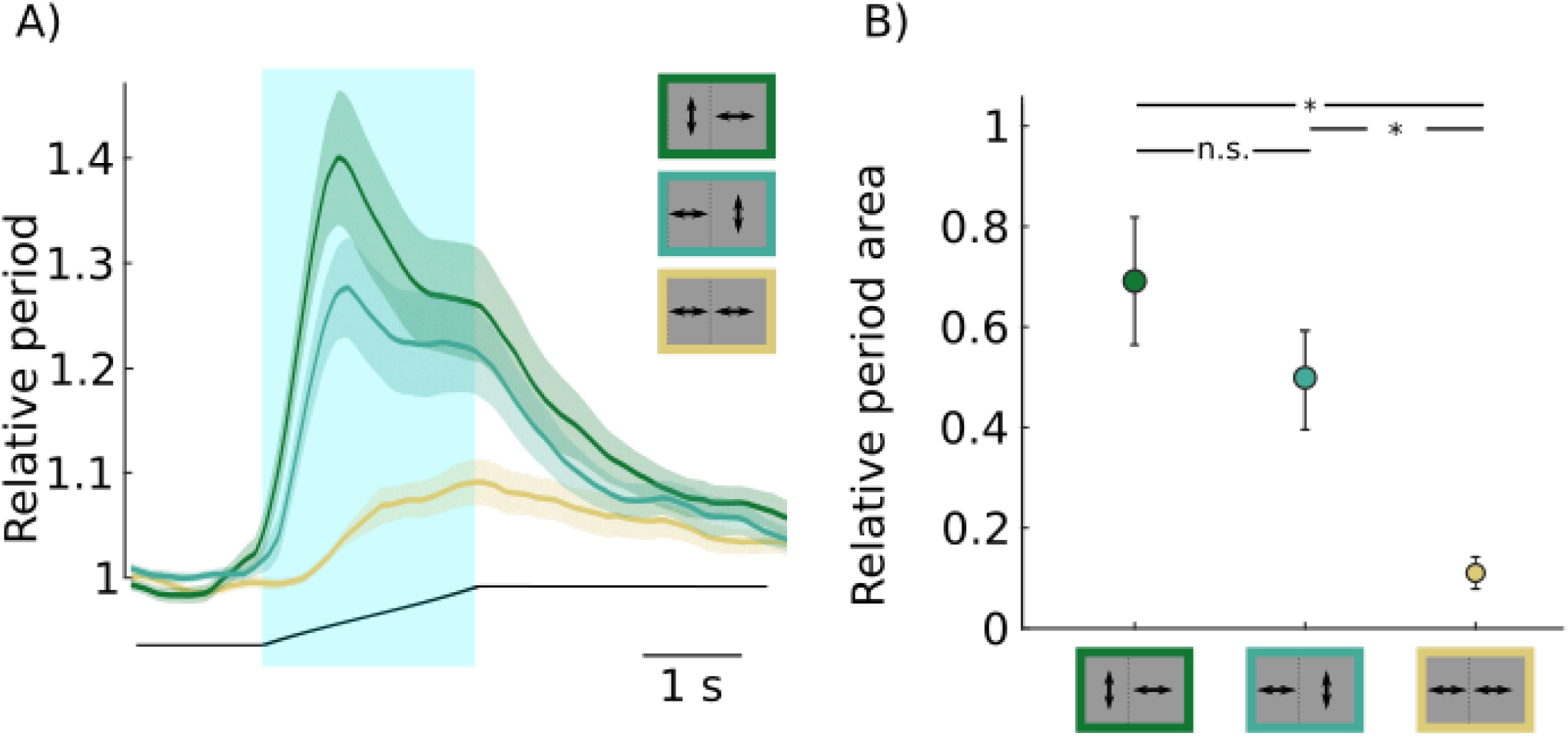
Cardiac responses to moving-edge stimuli with opposite polarization contrasts. (A) Animals’ relative cardiac period (mean (mean±s.e.m., N = 24) as a function of time in response to a moving edge. The light blue shaded area indicates the time of presentation of the moving edge. The black line below the plots illustrates the horizontal position of the moving edge in the screen. The stimuli consisted of a horizontally polarized edge moving over a vertically polarized background, a vertically polarized edge moving over a horizontally polarized background (greenish curves), and a horizontally polarized edge moving over a background of the same polarization but with a shift in the light-emitting lines (yellow curve; see Fig. S2B and table S2 for further stimuli details). (B) Relative period area integrated over the stimulation time (mean (mean±s.e.m., N = 24). Specific characteristics of the stimuli are detailed in table S2. Statistical differences are indicated as follows: n.s., non-significant differences; *, p < 0.01.

Although conspicuous behavioural responses to stimuli with polarization-only contrast have been reported in many invertebrate species, the relative salience of polarization contrast compared with intensity contrast has not been established. As a step in this direction, we compared the response to a polarization-only stimulus with the responses to stimuli of increasing intensity contrast (Fig. 4). We presented crabs with a vertically polarized edge moving over a horizontally polarized background of equal light intensity, and compared these responses with those evoked by dark edges moving over a light background (Michelson contrasts: 0.16, 0.31, 0.47, 0.64, 0.78, 0.91, 0.98, 0.99; N=35; further stimuli details in table S3). As previously reported, the cardiac response increases with the intensity contrast of the stimuli (Fig. 4B; Pérez-Schuster et al. (2023). To quantitatively compare the relative salience of the stimulus with polarization-only contrast with respect to the stimuli of intensity-only contrast, we used an inverse prediction. Specifically, we interpolated the mean response to the polarization-only stimulus onto the log-scale regression curve fitted to the intensity-contrast data (log(y) = b_0_ + b_1_*x; b_0_ = 0.86 *±* 0.02, b_1_ = 0.17 *±* 0.02; z_b1_ = 9.3, p_b1_<0.001). By means of a cluster bootstrap procedure (see Materials and Methods), we estimated that the magnitude of cardiac response to the polarization-only stimulus (cardiac response of VerticalE-HorizontalB = 0.461 *±* 0.003; N = 35) would be evoked by a stimulus of a Michelson contrast of 0.51 (95% CI= [0.31,0.74]). In other words, a polarization-only contrast stimulus like the one presented here has a saliency equivalent to a stimulus with an intensity contrast of 0.51.

**Figure 4.**
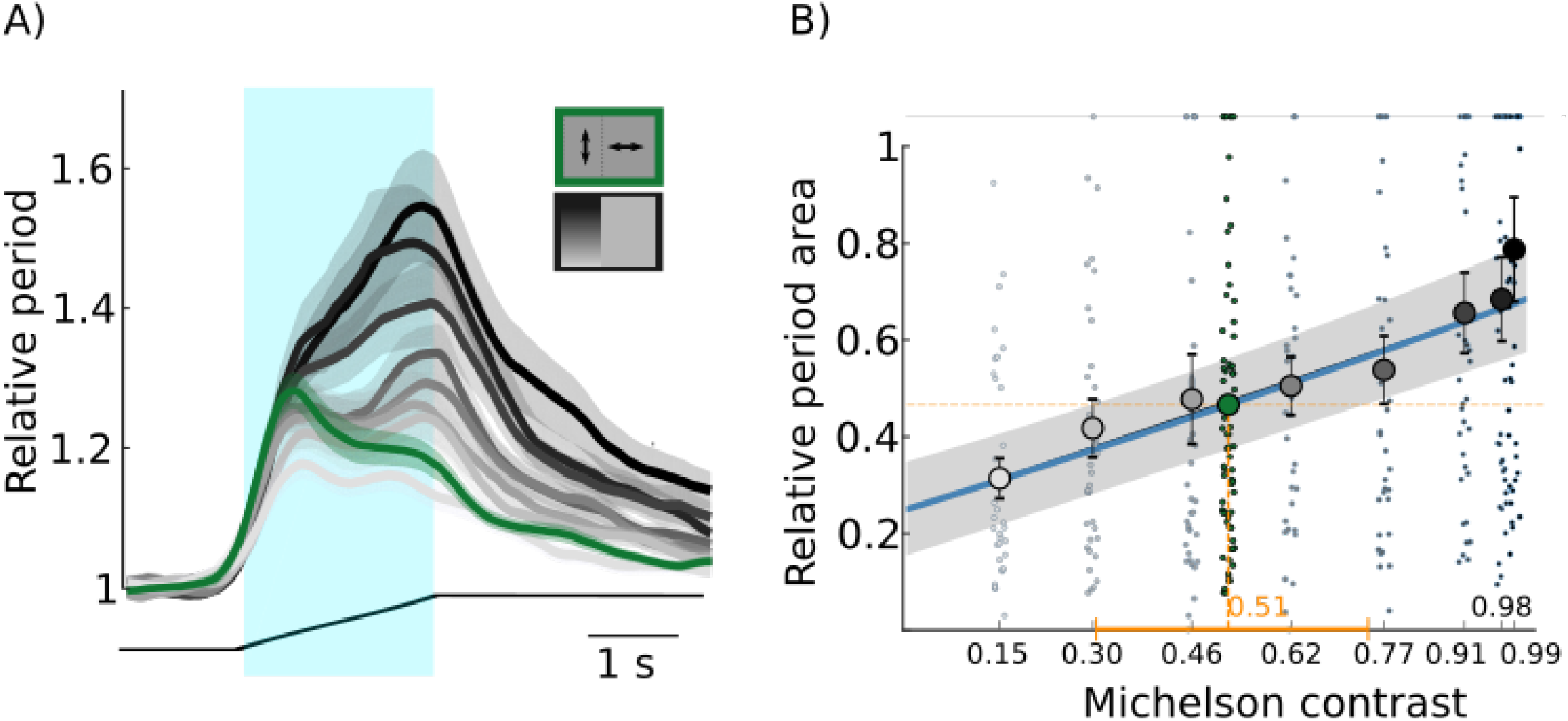
Cardiac responses of *Neohelice* to polarization-only and intensity-only-contrast stimuli. (A) Animals’ relative cardiac period (mean±s.e.m., N = 35) as a function of time in response to a moving edge. The light blue shaded area indicates the time of presentation of the moving edge. The black line below the plots illustrates the horizontal position of the moving edge in the screen. The stimuli consisted of intensity-contrast moving edges of increasing contrast (Michelson contrasts = 0.16, 0.31, 0.47, 0.64, 0.78, 0.91, 0.98, 0.99; POL contrast ≈ 0; increasing levels of grey in the figure represent higher contrasts), and a vertically polarized edge moving over a horizontally polarized background (Michelson contrast ≈ 0; POL contrast = 1.83; green curve). For further stimuli details see table S3. The upper inset summarizes the stimuli configuration. (B) Relative period area integrated over the stimulation time (mean±s.e.m., N = 35). The blue solid line and the grey shading represent the mean predicted values and their 95% confidence interval obtained from a log-scale regression curve fitted to the intensity-contrast data (*log*(*y*) = *b*_0_ + *b*_1_ *x*; *b*_0_ = 0.86±0.02, *b*_1_ = 0.17±0.02; z_*b*1_ = 9.3,p_*b*1_ *<* 0.001). Model predictions were back-transformed to the natural scale, and both predicted and observed cardiac period area values were normalized to the pre-stimulus average cardiac period area for visualization purposes only. The model itself was fitted to non-normalized data. The vertical orange dashed line indicates the mean intensity contrast equivalent in saliency to the polarization-only contrast. The horizontal orange solid line represents the 95% confidence interval [95% CI = 0.31, 0.74] of this intensity contrast. Specific stimuli characteristics are detailed in table S3.

Finally, we started studying how the perception of intensity contrast combines with that of polarization contrast. Specifically, we studied whether an intensity-contrast stimulus becomes more salient if it also possesses polarization contrast. Preliminary experiments showed no differences between the cardiac responses to high-intensity-contrast stimuli and those to stimuli combining the same intensity contrast with polarization contrast. Significant differences started to be detected when relatively low-intensity-contrast stimuli were combined with polarization contrast. Fig. 5 shows experiments in which the Michelson contrasts of the stimuli were 0.04 and 0.46 (upper and lower panels, respectively). The grey curves correspond to responses to intensity-only contrast stimuli, whereas the pink curves correspond to stimuli with the same intensity contrast combined with polarization contrast. For both intensity-contrast stimuli, the addition of polarization contrast significantly increased their salience (Fig. 5A; cardiac responses in the 0.46-intensity-contrast experiment: Int_contrast_= 0.46 *±* 0.03, Int_contrast_+POL_contrast_ = 0.56 *±* 0.03, Type II Wald *χ*^2^(1) = 5.7, p = 0.02, N = 23; Fig. 5B; cardiac responses in the 0.04-intensity-contrast experiment: Int_contrast_= 0.20 *±* 0.02, Int_contrast_+POL_contrast_ = 0.47 *±* 0.03, Type II Wald *χ*^2^(1) = 24.7, p<0.001, N = 25). Moreover, the effect of combining polarization with intensity contrast was more pronounced for the lower of the two intensity contrasts (differences in the cardiac responses in the 0.46-intensity-contrast experiment: (Int_contrast_+POL_contrast_)-(Int_contrast_) = 0.11 *±* 0.06; differences in the responses in the 0.04-intensity-contrast experiment: (Int_contrast_+POL_contrast_)-(Int_contrast_) = 0.28 *±* 0.05, Type II Wald *χ*^2^(1) = 4.8, p=0.03). Together, these results indicate that polarization contrast increases the salience of an intensity-contrast stimulus when its intensity contrast is low.

**Figure 5.**
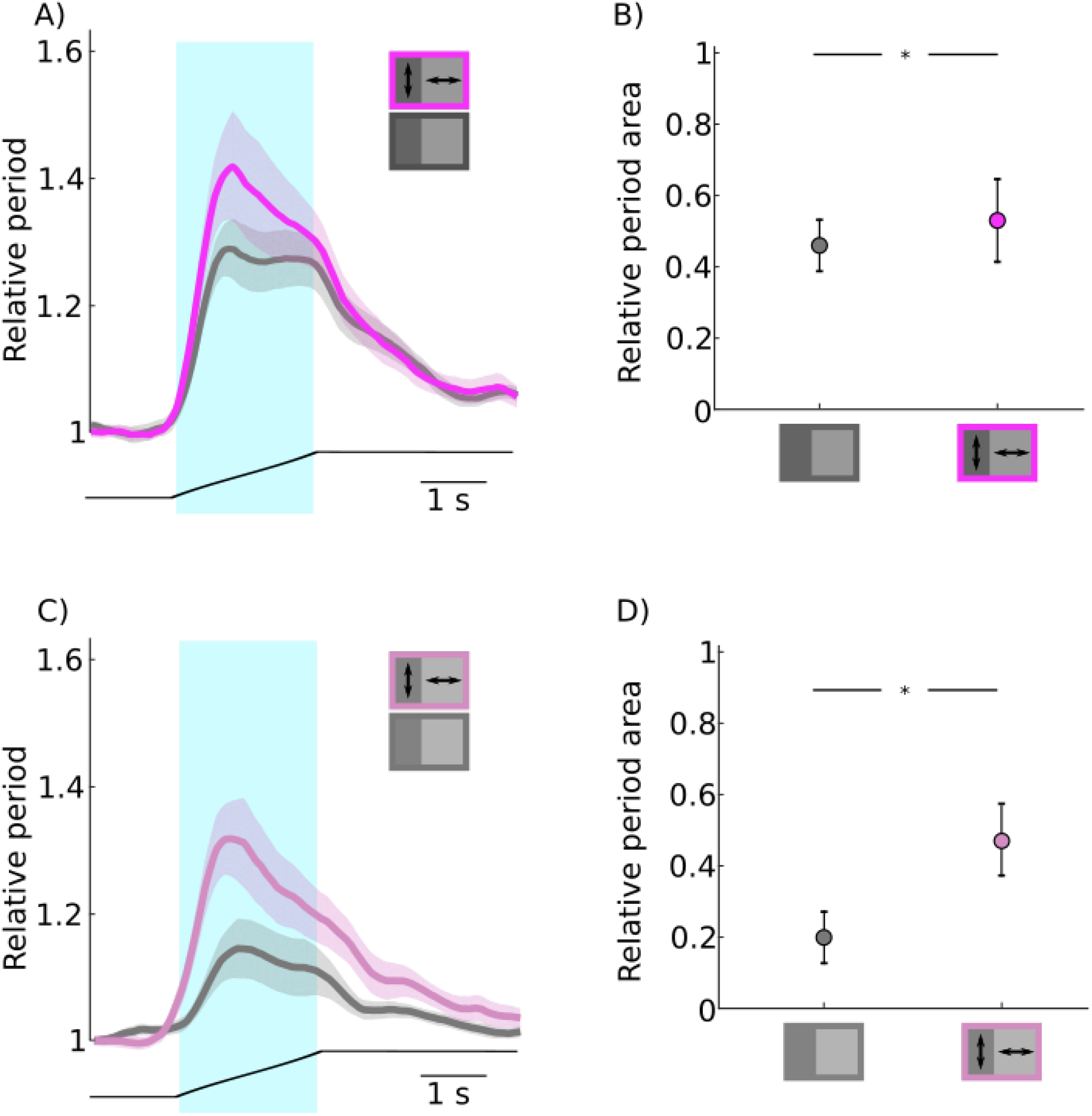
Interaction between polarization and intensity contrast in *Neohelice*. (A) Animals’ relative cardiac period (mean±s.e.m., N = 23) as a function of time in response to a moving edge. The stimuli consisted of a grey edge moving over a non-polarized light background (Michelson contrast = 0.46, POL contrast = 0.01; grey curve) or a stimulus with the same intensity contrast but with polarization contrast (Michelson contrast = 0.46, POL contrast = −1.82; pink curve). (B) Relative period area integrated over the stimulation time (mean±s.e.m., N = 23). (C) Animals’ relative cardiac period (mean±s.e.m., N = 25) as a function of time in response to the moving edge. The stimuli consisted of a grey edge moving over a non-polarized light background (Michelson contrast = 0.04, POL contrast = 0.01; grey curve) or a stimulus with the same intensity contrast but with polarization contrast (Michelson contrast = 0.04, POL contrast = −1.82; pink curve). (D) Animals’ relative cardiac period (mean±s.e.m., N = 23) as a function of time in response to the moving edge. For (A) and (B) the light blue shaded area indicates the time of presentation of the moving edge. Statistical differences: * stands for p < 0.05.

## DISCUSSION

Here we developed a visual stimulation device that allowed us to study the processing and integration of polarization and intensity contrast. Most recent studies on polarization vision have been carried out by presenting moving figures over backgrounds of different polarization but equal light intensity (e.g. R. M. Glantz and Schroeter (2007), Temple et al. (2012), M. How, Pignatelli, et al. (2012), and Basnak et al. (2018)). Thus, if the photoreceptor cells or downstream neurons (Alkaladi, M. How, and J. Zeil (2013) and Martín Berón de Astrada, Tuthill, and Tomsic (2009)) have different sensitivities to different polarizations of light, animal responses to these polarization-only stimuli could simply be the consequence of perceiving different light intensities between figure and background, and thereby detecting an intensity contrast (discussed in Labhart (2016)). Thus, we first studied whether there is a difference in sensitivity to vertically and horizontally polarized light in *Neohelice* (Basnak et al. (2018)). By presenting dark edges over vertically or horizontally polarized backgrounds, we found no difference in the animals’ cardiac response, suggesting that these backgrounds are perceived similarly by *Neohelice* (Fig. 2).

To further study whether *Neohelice* possesses different sensitivities to vertical and horizontal polarizations, we presented animals with two stimuli with polarization contrast but no intensity contrast: a vertically polarized edge moving over a horizontally polarized background, and the opposite combination (Fig. 3). The responses to these stimuli were highly similar. However, we had previously observed that stimuli of opposite intensity contrasts evoke largely different responses (Bengochea (2017)). Like other animal species, *Neohelice* responds very weakly to stimuli in which a light figure moves over a dark background (e.g. Santer, Simmons, and Rind (2005) and Yilmaz and Meister (2013)). Thus, in contrast to the strong asymmetry observed in the responses to intensity-contrast stimuli of opposite polarity, cardiac responses to opposite polarization contrasts were highly similar, suggesting that polarization sensitivity involves processing mechanisms distinct from the differential perception of light intensity between vertically and horizontally polarized light. Overall, these results are in line with the idea that *Neohelice* processes polarization contrast in a channel different from that processing intensity contrast.

Our next experiment aimed to obtain a precise estimation of the relative salience of a polarization-only contrast stimulus with respect to an intensity-only contrast stimulus under our laboratory conditions (Fig. 4. By measuring the area under the curve of the cardiac response over the entire visual stimulation period, we estimated that a polarization-only contrast stimulus has a salience equivalent to an intensity-only contrast stimulus of 0.51 on the Michelson contrast scale. While obtaining an estimate of the relative salience of polarizationand intensity-contrast stimuli was the simple goal of this experiment, we observed that the temporal profile of the cardiac response to stimuli of equivalent salience differed between polarization and intensity stimuli. For the polarization stimulus, the response peaked close to the beginning of visual stimulation, whereas for the intensity stimulus the peak response occurred towards the end of stimulation. We characterised in depth the temporal profile of the cardiac response of *Neohelice* when confronted to the same visual stimuli used here (Pérez-Schuster et al. (2023)). Briefly, the cardiac response is composed of two sequential phases. The peak of the first phase increases with stimulus contrast within a low-contrast range and saturates for high-contrast stimuli. In turn, the second phase appears at high contrasts, becoming more pronounced as contrast increases. Here, we find that, for a polarizationcontrast stimulus, the second-phase response is lower than that expected for an equivalent intensity-contrast stimulus (Fig. 4A). This might seem a subtle difference, however, throughout every experiment in the current study in which a polarization-only contrast stimulus was presented, both the magnitude and the temporal profile of the response were highly similar, which indicates that the recorded response profile to polarization-only stimuli is highly consistent.

Although we have no explanation for the difference in the temporal profile of the cardiac response between polarization- and intensity-contrast stimuli, this difference suggests once again that these stimuli are likely processed through distinct contrast channels. Under this hypothesis, the polarization contrast channel would strongly feed the neural circuits that trigger the first phase of the cardiac response, but would not feed, or would weakly feed, those triggering the second phase. Alternative explanations, however, remain possible. Here, we compared the responses to a polarization-only stimulus with the responses to intensity-contrast stimuli in which the edge was darker than the background. We did not perform a systematic study of cardiac responses when intensity-contrast stimuli have the opposite contrast, i.e. a light edge moving over a dark background. Preliminary results in this direction suggest that a high-intensity-contrast stimulus of such polarity evokes a cardiac response of similar magnitude and temporal profile to that evoked by polarization-only contrast stimuli (Salomon (2020)). This raises the possibility that a polarization-only contrast stimulus could be perceived as a stimulus in which the edge is lighter than the background. In our stimuli, the presentation of the background for 10 minutes before the appearance of the edge could induce an adaptation of the polarization channel that matches that of the background. Then, when the edge appears with the same intensity as the background but with an orthogonal polarization that mainly stimulates the non-adapted channel, this could be sensed as a general increase in light intensity. Although this hypothesis is rather speculative, it is at the same time reasonable, and highlights that further experiments are needed to better understand the integration of polarization and intensity information in polarization-sensitive animals.

Finally, we studied whether the combination of polarization and intensity contrast cues enhances target detection (Fig. 5). To this end, we compared stimuli combining polarization and intensity contrast with stimuli that presented only intensity contrast. Preliminary experiments showed that the addition of polarization cues to high-intensity-contrast stimuli produced no enhancement of the animals’ responses. Conversely, stimuli combining polarization and low-intensity contrast increased the animals’ response relative to the stimuli presenting only intensity contrast (Fig. 5), indicating that this particular combination of contrast cues effectively produced an enhancement in contrast detection. These results suggest that, as long as intensity-contrast information is highly salient, the animals rely on this cue to guide behaviour, and that they combine it with polarization information in more ambiguous situations, where intensity contrast is low. Our results are consistent with the principle of inverse effectiveness, which states that a response enhancement produced by combining different stimuli is greatest when the individual stimuli are weakly effective on their own (Stein et al. (2009)). Recently, a study performed in fiddler crabs by Smithers (2019) challenges the classical notion of contrast enhancement, the idea that polarization cues reinforce or amplify intensity-contrast detection. The authors propose that these crabs process polarization and intensity-contrast information independently and in parallel, and that their response to visual stimulation follows whichever contrast is more salient, independently of the less salient one. The authors conclude that this strategy broadens the range of detectable information, increasing the likelihood of detecting moving objects. Our results can be explained in terms of the mechanisms proposed in this study. However, to evaluate this hypothesis rigorously in *Neohelice*, future experimental designs should include not only an intensity-only contrast stimulus and a stimulus combining polarization and intensity contrast, but also a polarization-only contrast stimulus.

Vision has evolved in relation to the particular environmental conditions and behavioural requirements of each species. Many fiddler crab species and *Neohelice* inhabit mudflats, a habitat that is relatively simple in terms of visual information content and highly spatially structured in two dimensions (Jochen Zeil and Hofmann (2001) and J. Zeil (2023)). Their compound eyes show specific adaptations to the visual conditions of an essentially flat world (J. Zeil and Al-Mutairi (1996) and M. Berón de Astrada et al. (2012)). The eyes cover the entire 360° panorama and possess an equatorial acute zone, aligned with the visual horizon, in which vertical resolving power is greatly enhanced. The horizon divides the two principal sources of polarized light in the mudflat: sky polarization produced by scattering, and ground polarization produced by reflections from the mudflat surface (J. Zeil (2023)). Polarization vision is expected to depend on how the structure of the polarized scenes are organised in the environment. Thus, the particularities of sensing and processing polarization information arising from the sky or from ground reflections are also expected to be regionalised (Sabra and R. M. Glantz (1985)). So far, we have treated the processing of polarization-contrast information as if a single, general mechanism underlay the processing of this visual attribute. However, future studies on polarization vision should consider that the processing of polarization information may differ depending on the region of the visual scene involved.

## MATERIALS AND METHODS

### Animals

Male *Neohelice granulata* crabs were collected from the rías (narrow coastal inlets) of San Clemente del Tuyú, Argentina. Only adult male animals 2.7 - 3.0 cm width across the carapace were included in the current study. The animals were transported to the laboratory where they were kept in plastic tanks filled with up to 1 cm tall of artificial marine water (Red Sea’s Coral Pro Salt; salinity 10 14%, pH 7.4 - 7.6). In the laboratory animals were kept on a 12 h light/dark cycle (lights on 7:00 AM to 7:00 PM). All the experiments were run during the light cycle. The experimental protocols were performed in accordance with relevant guidelines and ethical regulations of the School of Science, Universidad de Buenos Aires.

### Electrocardiogram recordings

Across many invertebrate and vertebrate species sensory stimuli evoke changes in cardiac activity (Cuadras (1980), Laming and Austin (1981), Ide and Hoffmann (2002), and Barrios, Farias, and Moita (2021)). The intensity of these changes have been used as an indicator of perceptual sensitivity (e.g. Schenberg, Vasquez, and Costa (1993)). In crabs, sensory stimuli of different modalities evoke a transient reduction in heart rate (Burnovicz, Oliva, and Hermitte (2009) and Pérez-Schuster et al. (2023)). In *Neohelice*, we have observed that the intensity of the cardiac response increases with both intensity- and polarization-only contrast stimuli (Pérez-Schuster et al. (2023) and Basnak et al. (2018)). We have found that this response presents several advantages with respect to other behavioral variables for estimating visual perception, especially when measuring responses to stimuli of low or moderate salience, as is the case here. One advantage is that this is a highly sensitive measure that presents little error. In addition, we have observed that many animals do not change their behavior when confronted with stimuli of low salience; however, they develop small but highly robust changes in their heart rate, which indicate that they perceive them (Burnovicz, Oliva, and Hermitte (2009) and Pérez-Schuster et al. (2023)).

The methodology for performing the electrocardiogram recordings has been explained in detail in Hermitte and Maldonado (2006). Briefly, we used silver wire electrodes (diameter 0.25 mm, A-M Systems, Carlsborg, USA) to monitor heart rate. A small jack with two metallic pins, to which the electrodes were soldered, was cemented to the dorsal carapace of the crabs above the heart. The free ends of the electrodes were inserted 2-4 mm into holes drilled 4-5 mm apart in the cardiac region of the dorsal carapace and secured with super glue. An impedance converter (model 2991, UFI, California, USA) connected to the metallics pins was used to monitor the heart rate. The output signal of the impedance converter allowed monitoring of the cardiac activity as a measure of dynamic resistance (Yang, Carbó Tano, and Hermitte (2013)). The converter’s output was digitized using Digidata 1440A (Molecular Devices, Sunnyvale, CA, USA). All experiments were performed at least 2 days after attaching the jack and electrodes.

### Experimental arena

The experimental arena consisted of a large box with four white foam-board walls, 100 *×* 70 cm each, placed at 45° from the vertical (Fig. 1). The flanking walls were oriented in this way to avoid differential reflection of vertically and horizontally polarized light (Basnak et al. (2018)). The monitor screen was placed at one end of the box and the other end presented a white foam cover. To perform the electrocardiogram recordings, the animal was tethered by securing its claws and legs against its body with a rubber band. It was then positioned 20 cm from the center of the monitor screen and held in place with an adjustable clamp; finally, the impedance converter was plugged into the jack.

### Intensity and polarization screen

We had found that *Neohelice* possesses an orthogonal receptor arrangement for the detection of polarized light in which two polarization channels are aligned with the vertical and horizontal orientations (Basnak et al. (2018)). To deliver polarized visual stimuli tuned to these polarization channels while controlling color, we modified a commercial 3D LCD monitor display (Fig. S1A; LG-32LA613B, LG Electronics, Seoul, South Korea). This screen emits differently polarized light from odd and even horizontal lines of pixels. Human eyes can hardly discriminate between horizontal lines of pixels in commercial screens, even at relatively close distance to the monitor. Thus, we reasoned that arthropod eyes, which possess much lower sampling resolution, would perceive light coming from pixels in neighboring horizontal lines as coming from the same point in space. That is, a phenomenon similar to what happens to us when we watch a color monitor: we do not discriminate the separation between neighboring color pixels, and we perceive light with different spectral compositions as coming from the same direction in space. Particularly, the monitor screen we modified emits right-elliptically polarized light for pixels located in even rows and left-elliptically polarized light for those in odd rows (refresh rate 60 Hz). By placing and properly orienting an achromatic quarter-wave retarder sheet just in front of the screen, it is possible to obtain vertically (90°) and horizontally (0°) linearly polarized light coming from odd and even rows, respectively Fig. S1A. To this end, an achromatic quarter-wave retarder sheet (30 *×* 30 cm, 450–700 nm, #88-253, Edmund Optics, Fig. S1B) was mounted on a rectangular window cut into a black cardboard sheet. The sheet mounted on the cardboard was located directly over the screen in order to minimize light crossing the retarder diagonally. The window was 18 cm wide and 19 cm high and as the crab was located 20 cm away from the screen, the stimulation area encompassed 48° *×* 52°. From this distance the angular separation between two horizontal rows of pixels was 0.14°. The crabs were positioned with the lateral pole of one of their eyes looking at the center of the stimulation window. As other crabs, *Neohelice*, possess high panoramic compound eyes with a narrow horizontal band (10–20°) of high vertical resolution (M. Berón de Astrada et al. (2012)). In the lateral pole of the eye, the minimum vertical and horizontal interommatidial angles are 0.6° and 0.9° respectively (M. Berón de Astrada et al. (2012), Fig. S2A). Thus, if we assume that the acceptance angle of the ommatidia matches the interommatidial angle, theoretically more than 4 horizontal and 6 vertical lines are fused at the peak sampling region of the eye (Fig. S2A). For ommatidia outside the horizontal fovea, the number of fused lines per ommatidia is much larger.

Typically, by controlling the RGB values of the pixels on a screen, it is possible to control the intensity of the light emitted. Here we always used gray emissions, i.e. the RGB values for the red, green and blue pixels were the same. To calculate the intensity of the light cast by the screen, we placed a light guide connected to a spectrophotometer (OceanOptics USB2000) at the position where crabs were located within the experimental arena pointing toward the center of the screen.

Remarkably, with our modified screen, by differentially controlling the RGB values of the odd and even pixels, it was possible to control the proportion of vertically to horizontally linearly polarized light cast by the screen. Vertically polarized light (I_v_) was measured as the light intensity emitted by the monitor screen and transmitted through a vertically oriented linear polarizer, and horizontally polarized light (I_h_) as the light intensity transmitted through a horizontally oriented one. We define the 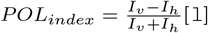, as the difference between vertically and horizontally polarized light intensities, relative to the total intensity of light emitted. Theoretically, the POLindex (POLI) ranges from +1 (fully vertically polarized light) to −1 (fully horizontally polarized light), with 0 indicating equal intensities at 0° and 90°. In practice, light emitted with only odd rows on (RGB = 255) was mainly vertically polarized at 88.2°, with a POL_index_ of 0.93 (table S2; Fig. S1B), while light emitted with only even rows on (RGB = 255) was mainly horizontally polarized at 1.6°, with a POL_index_ of −0.90 (table S2; Fig. S1B).

### Visual stimuli

The visual stimulus consisted of a vertical edge translating horizontally across the screen displaying a uniform background (Pérez-Schuster et al. (2023)). This simple stimulus consistently evokes defensive responses in crabs (Bengochea (2017) and Pérez-Schuster et al. (2023)). At the border of the advancing edge, the following transition occurred in the columns of pixels on the screen: as the background pixels turned on or off, the edge pixels turned on or off at the same moment. The temporal on/off progression of the border across the screen defined the edge’s velocity. This stimulus has the advantage that, by presenting a single contrast transition, the effective stimulus contrast can be easily determined (Nordström, Bolzon, and O’Carroll (2012)). In the current study, the edge occupied the entire height of the stimulation area and advanced from the anterior-to the posterior-lateral visual field of the animals (M. Berón de Astrada et al. (2012)) at a constant angular speed of 30° s^-1^ (Pérez-Schuster et al. (2023)).

Throughout current study we presented different combinations of edges translating over different backgrounds. Each edge or background is characterized by a particular polarization and light intensity. We will call this edge/background combination a stimulus. Supplementary tables S1 to S5 detail the characteristics of the stimuli presented in the different experiments of current study and how we produced them. The intensity contrast of a stimulus was calculated with Michelson’s equation (Michelson (1927)). In our two-channel polarization system, polarization contrast (POL_contrast_) was estimated as the difference between the POL_index_ of the edge and that of the background, 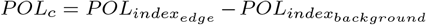 [2]. Thus, theoretically the POL contrast of the stimuli range from +2 (a fully vertically polarized edge advancing over a fully horizontally polarized background) to −2 (opposite polarization of edge and background), with 0 indicating no difference between the polarization indices of the edge and background.

To confirm that crabs fuse light arriving from neighboring lines of the screen and, therefore, that we could modulate stimulus polarization contrast by turning neighboring lines on or off while still controlling the luminance of the stimulus, we performed the control experiment shown in Fig. S2. In this experiment, we displayed a horizontally polarized background by turning on lines 2, 6, 10, 14, etc. (RGB = 255). In one of the experimental stimuli, over this background we presented a polarized edge of the same intensity, also horizontally polarized, but produced by turning on lines 4, 8, 12, 16, etc. (RGB = 255) and off lines 2, 6, 10, 14, etc. (RGB = 0) (illustrated in the right panel of Fig. S2B and quantified in table S5). This is a stimulus that presents a markedly different spatial light pattern between edge and background when viewed through a high-resolution imaging system, and would be expected to elicit some response if the animals could discriminate the differences. However, the cardiac activity of the animals did not change when confronted with this stimulus (group highlighted with blue color in Figs. S2C and S2D), indicating that animals merge the light coming from neighboring lines of pixels. In addition, also keeping edge and background intensity equal, we presented different stimuli in which we progressively increased the proportion of vertically polarized light of the edge by differentially turning on even rows (3, 7, 11, 15, etc.) and odd rows (4, 8, 12, 16, etc.). In this way, we kept the Michelson contrast close to 0 while increasing the polarization contrast. As expected, the more content of vertically polarized light of the edge (i.e. the more the polarization contrasts with the horizontally polarized background) the greater the cardiac response of the animals (Figs. S2C and S2D; Basnak et al. (2018)).

### Experimental protocol and data analysis

Each crab was placed in the experimental arena and left undisturbed for 10 minutes until the first stimulus presentation. Thirty seconds after the moving stimulus reached the end of the screen, the background illumination of the following stimulus was displayed. The intertrial interval was 10 min and the order of stimulus presentation to each animal was randomized. The synchronization between visual stimulation and cardiac activity recording was implemented using Matlab (MathWorks, Natick, MA, USA) and the Psychophysics Toolbox extensions (D.H. Brainard (1997), D.G. Pelli (1997), and Kleiner, D. Brainard, and D. Pelli (2007)). Each of the experiments presented in this work has an associated supplementary table where we describe the main characteristics of the stimuli. The cardiac responses to intensity-contrast stimuli shown in Fig. 4 were previously reported in Pérez-Schuster et al. (2023). In the present work, we reanalyzed these data together with the results from an experimental group stimulated with a polarization-only stimulus (Fig. 4B).

To quantify the cardiac activity, we applied the same methodology used in previous studies from our group (for more details see Pérez-Schuster et al. (2023)). Briefly, we measured the period between two beats and normalized it to the average period during the 10 s prior to stimulation. For analytical purposes, we assigned the relative period value to a time point equidistant between the two beats. Then, we linearly interpolated the data between every two consecutive relative periods for each ECG recording. After interpolation, we obtained a continuous temporal profile of the cardiac activity for each stimulation trial performed on each animal. Finally, we integrated the area under the temporal profile curve during stimulus presentation to quantify the cardiac response to visual stimulation. This methodology allowed a temporal and quantitative analysis of the cardiac response across large numbers of individuals.

### Statistics analysis

All analyses were conducted in R (v4.4.3). Either general linear mixed models in log scale or generalized linear mixed models (GLMM) with Gamma distribution (log link function) were fitted to model the variations in the area under the temporal curve of cardiac period during stimulation time. Both statistical approximations are practically equivalent in order to model continuous variables with asymmetrical distributions and were chosen in any case based on the Akaike information criterion (AIC). Models were run using the nlme and glmmtmb packages. The type of visual stimulus displayed to the animal was used as a categorical fixed effect in experiments of Figs. 2, 3 and 5, including four, three and two levels, respectively. Crab identity was specified as a random effect since each one was presented with all types of stimuli in every experiment. If the fixed effect was found to be significant, Tukey post-hoc contrasts were conducted using the emmeans package to assess pairwise comparisons between types of stimuli. Model assumptions were evaluated using the DHARMa package.

In the experiment of Fig. 4, the fixed effect was the intensity contrast of the stimulus and was included in the model not as a categorical but as a continuous variable in order to fit a linear regression analysis in log scale of the response variable. Then, an inverse interpolation was performed with data of trials where a polarization contrast stimulus was displayed to the animals, with the aim of obtaining the equivalent intensity contrast that can generate the responses to polarization. Briefly, the process consisted of a cluster (crab-level) bootstrap procedure to propagate uncertainty from both the regression and the polarization-contrast response sample into the estimated equivalent intensity contrast. In each of 2000 bootstrap iterations, crabs were resampled with replacement (separately for the intensity-contrast dataset and the polarization-contrast dataset), preserving all observations belonging to each resampled individual to maintain the within-animal correlation structure. The general linear mixed model relating area under the curve to intensity contrast was refitted on each resampled dataset, yielding bootstrap estimates of the intercept and slope. The mean response to the polarization-contrast stimulus was recalculated on the corresponding resampled polarization dataset, and the equivalent intensity contrast was obtained by algebraically inverting the fitted regression line (i.e., solving for the contrast value predicted to produce the observed polarization response). This procedure generated an empirical bootstrap distribution of 2,000 equivalent intensity contrast estimates, from which a point estimate and 95% confidence interval (2.5th and 97.5th percentiles) were derived.

## ACKNOWLEDGMENTS

This research was conducted in the context of a severe funding crisis affecting Argentine science. The crisis is marked by budget cuts, salary stagnation, and reduced support for public research institutions. This study was partially supported by the Argentinean National Research Agency (ANPCYT) PICT A-2018-01677 to M.B.A and PICT 2018-1672 and PICT 2017-0251 to V.P.S.. Despite these adverse conditions, we thank the Facultad de Ciencias Exactas y Naturales, University of Buenos Aires, and CONICET for providing the minimal conditions needed to carry out this study. We also thank our team and iB3 for their support.

## SUPPLEMENTARY FIGURES

**Figure S1.**
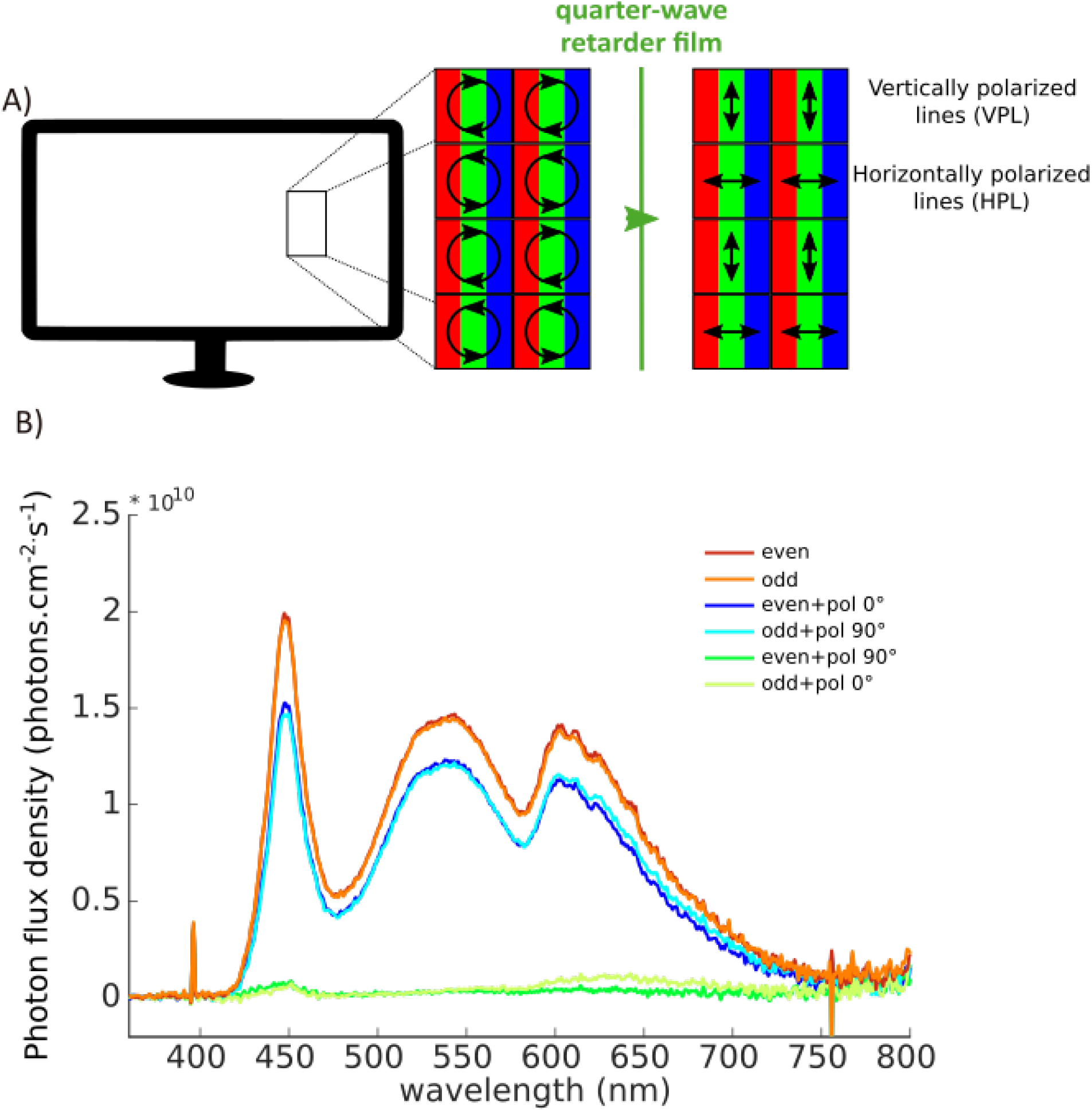
Experimental screen. (A) Our experimental 3D display emits circularly polarized light that becomes vertically (VPL, odd lines) or horizontally (HPL, even lines) polarized after passing through a quarter retarder lm. (B) Spectral intensity of the light (photon flux density) emitted by the monitor screen as a function of wavelength, under different conditions. The even and odd lines, when on at an RGB value of 255, emit highly similar intensities across all wavelengths (red and orange curves). Spectral recordings for the blue and light-blue curves were performed by placing a linear polarizer between the monitor and the spectrophotometer, with its transmission axis parallel to the angle of polarization of the corresponding emitting lines (blue = even lines on; light-blue = odd lines on). Conversely, spectral recordings for the green and light-green curves were performed by placing a linear polarizer between the monitor and the spectrophotometer, with its transmission axis perpendicular to the angle of polarization of the corresponding emitting lines (green = even lines on; light-green = odd lines on).

**Figure S2.**
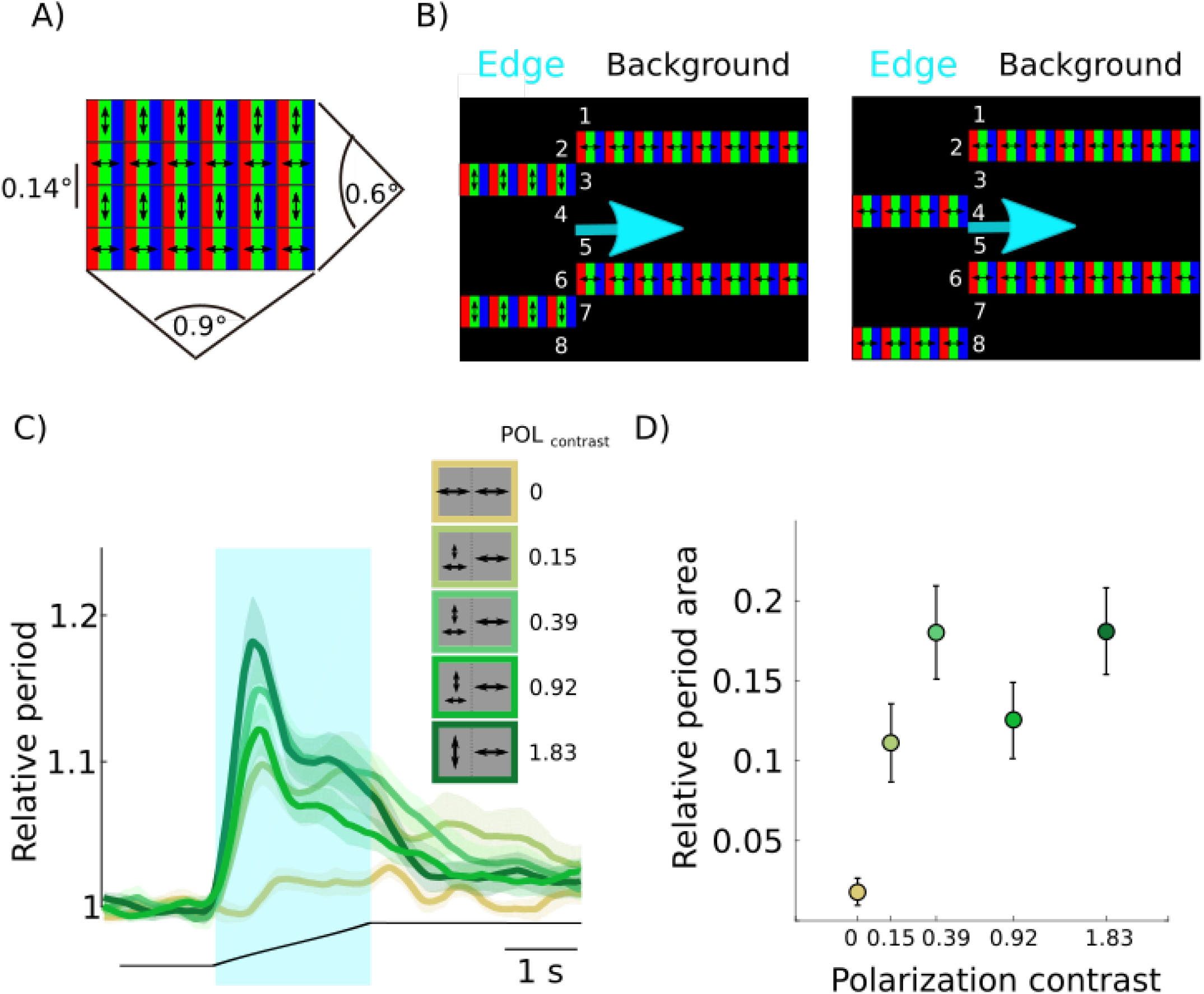
Control of the optical fusion of the interlaced monitor lines. (A) From the crab’s position, 20 cm away from the screen, the minimum angular separation between horizontal lines is 0.14°. In the lateral pole of the eye, the minimum vertical and horizontal interommatidial angles are 0.6° and 0.9° respectively (M. Berón de Astrada et al. (2012)). Thus, if the acceptance angle matches the interommatidial angle, around four horizontal and six vertical monitor lines are fused at the eye equator, the region of highest sampling frequency in the lateral part of the eye. (B) Stimuli used to control for the spatial disparity between the edge and background light patterns. Left panel: a vertically polarized edge over a horizontally polarised background (Michelson contrast 0; POLcontrast = 1.83). Right panel: a horizontally polarized edge over a horizontally polarized background (Michelson contrast 0; POL_contrast_ = 0). (C) Animals’ relative cardiac period (mean±s.e.m., N = 36) as a function of time in response to a moving edge. The light blue shaded area indicates the time of presentation of the moving edge. The black line below the plots illustrates the horizontal position of the moving edge in the screen. The stimuli consisted of edges of increasing polarisation contrast (POL_contrast_) moving over a horizontally polarised background (Michelson contrasts 0; POL_contrast_ = 0, 0.15, 0.39, 0.92, 1.83) (see **??** and table S5 for further stimuli details). Note that the stimulus with the highest spatial disparity between edge and background (right panel in B; Michelson contrast 0; POL_contrast_ = 0) is the one that shows the minimum response, suggesting that the monitor lines are being fused in the ommatidial array. (D) Relative period area integrated over the stimulation time (mean±s.e.m., N = 36). Note that the cardiac response increases with polarization contrast. The x-axis is displayed on a logarithmic scale.

## SUPPLEMENTARY TABLES

**Table S1.**
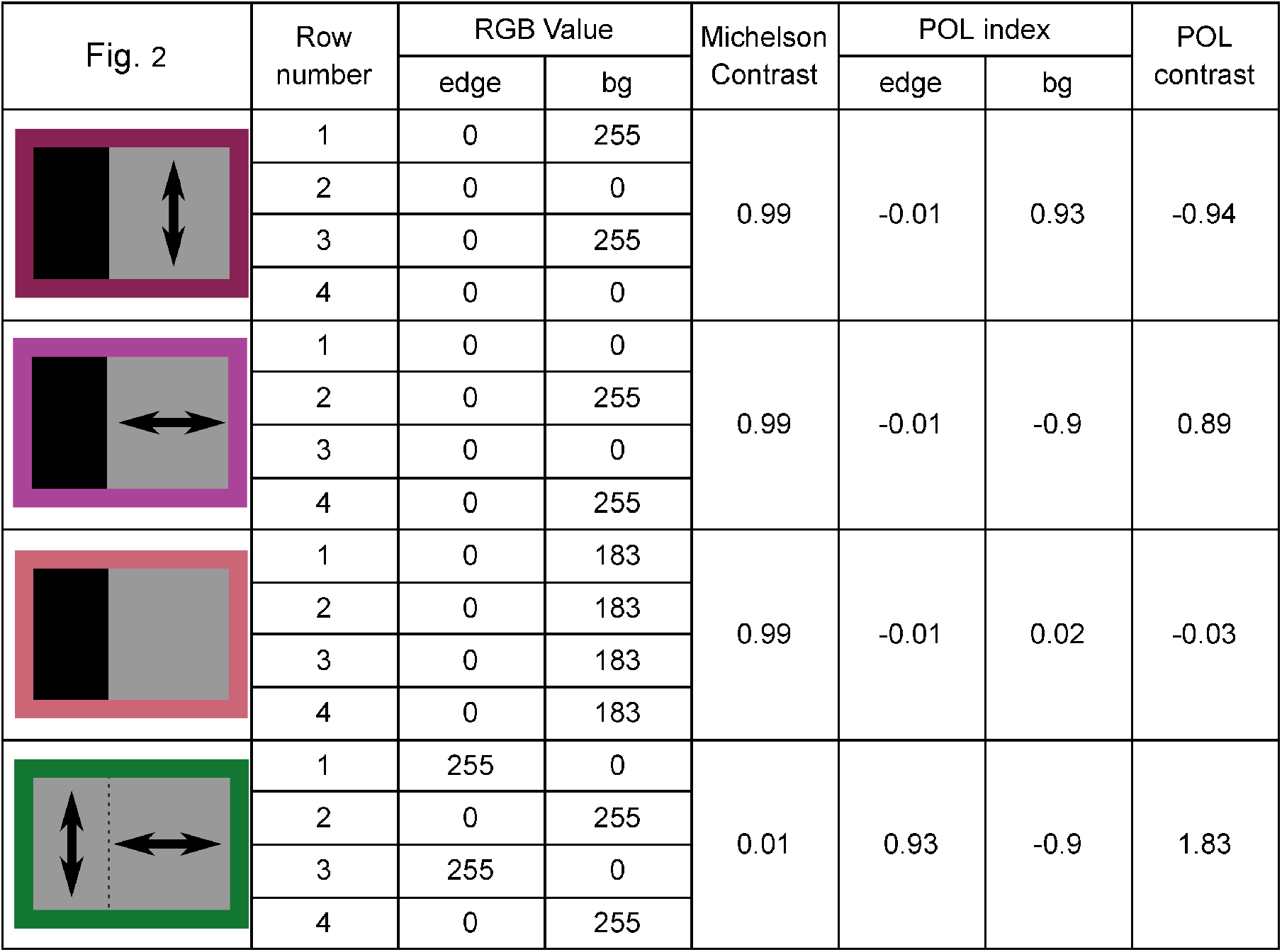
Stimuli characteristics for the experiment of Fig. 2. For each experimental condition, the RGB value of each row, the Michelson contrast between edge and background, the Polarization index (see Intensity and polarization screen), and the Polarization contrast (see Visual stimuli) are specified. The abbreviation bg stands for background.

| Fig. 2 | Row number | RGB Value |  | Michelson Contrast | POL index |  | POL contrast |
| --- | --- | --- | --- | --- | --- | --- | --- |
|  |  | edge | bg |  | edge | bg |  |
|  | 1 | 0 | 255 | 0.99 | -0.01 | 0.93 | -0.94 |
|  | 2 | 0 | 0 |  |  |  |  |
|  | 3 | 0 | 255 |  |  |  |  |
|  | 4 | 0 | 0 |  |  |  |  |
|  | 1 | 0 | 0 | 0.99 | -0.01 | -0.9 | 0.89 |
|  | 2 | 0 | 255 |  |  |  |  |
|  | 3 | 0 | 0 |  |  |  |  |
|  | 4 | 0 | 255 |  |  |  |  |
|  | 1 | 0 | 183 | 0.99 | -0.01 | 0.02 | -0.03 |
|  | 2 | 0 | 183 |  |  |  |  |
|  | 3 | 0 | 183 |  |  |  |  |
|  | 4 | 0 | 183 |  |  |  |  |
|  | 1 | 255 | 0 | 0.01 | 0.93 | -0.9 | 1.83 |
|  | 2 | 0 | 255 |  |  |  |  |
|  | 3 | 255 | 0 |  |  |  |  |
|  | 4 | 0 | 255 |  |  |  |  |

**Table S2.**
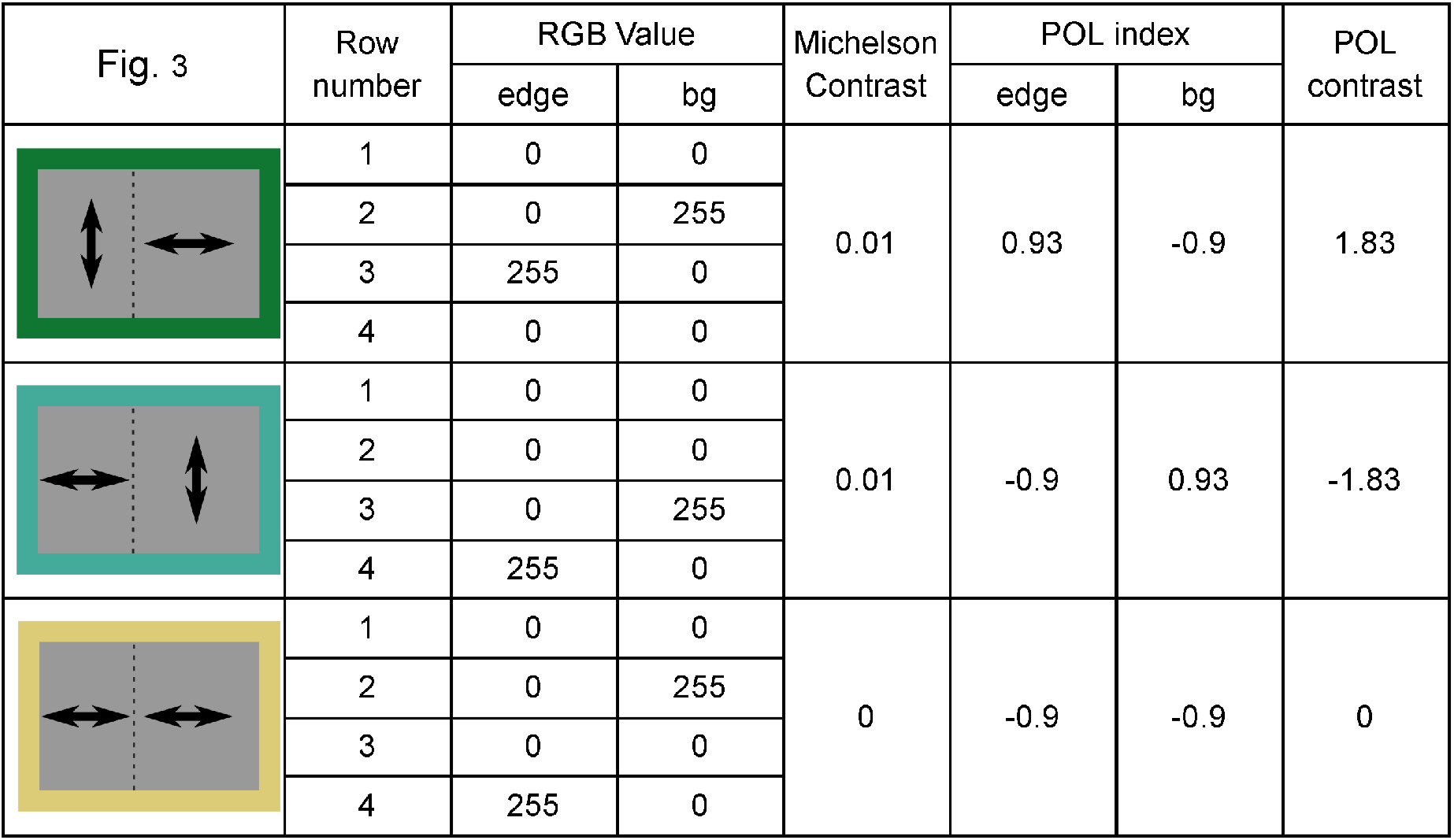
Stimuli characteristics for the experiment of Fig. 3. For each experimental condition, the RGB value of each row, the Michelson contrast between edge and background, the Polarization index (see Intensity and polarization screen), and the Polarization contrast (see Visual stimuli) are specified. The abbreviation bg stands for background.

| Fig. 3 | Row number | RGB Value |  | Michelson Contrast | POL index |  | POL contrast |
| --- | --- | --- | --- | --- | --- | --- | --- |
|  |  | edge | bg |  | edge | bg |  |
|  | 1 | 0 | 0 | 0.01 | 0.93 | -0.9 | 1.83 |
|  | 2 | 0 | 255 |  |  |  |  |
|  | 3 | 255 | 0 |  |  |  |  |
|  | 4 | 0 | 0 |  |  |  |  |
|  | 1 | 0 | 0 | 0.01 | -0.9 | 0.93 | -1.83 |
|  | 2 | 0 | 0 |  |  |  |  |
|  | 3 | 0 | 255 |  |  |  |  |
|  | 4 | 255 | 0 |  |  |  |  |
|  | 1 | 0 | 0 | 0 | -0.9 | -0.9 | 0 |
|  | 2 | 0 | 255 |  |  |  |  |
|  | 3 | 0 | 0 |  |  |  |  |
|  | 4 | 255 | 0 |  |  |  |  |

**Table S3.**
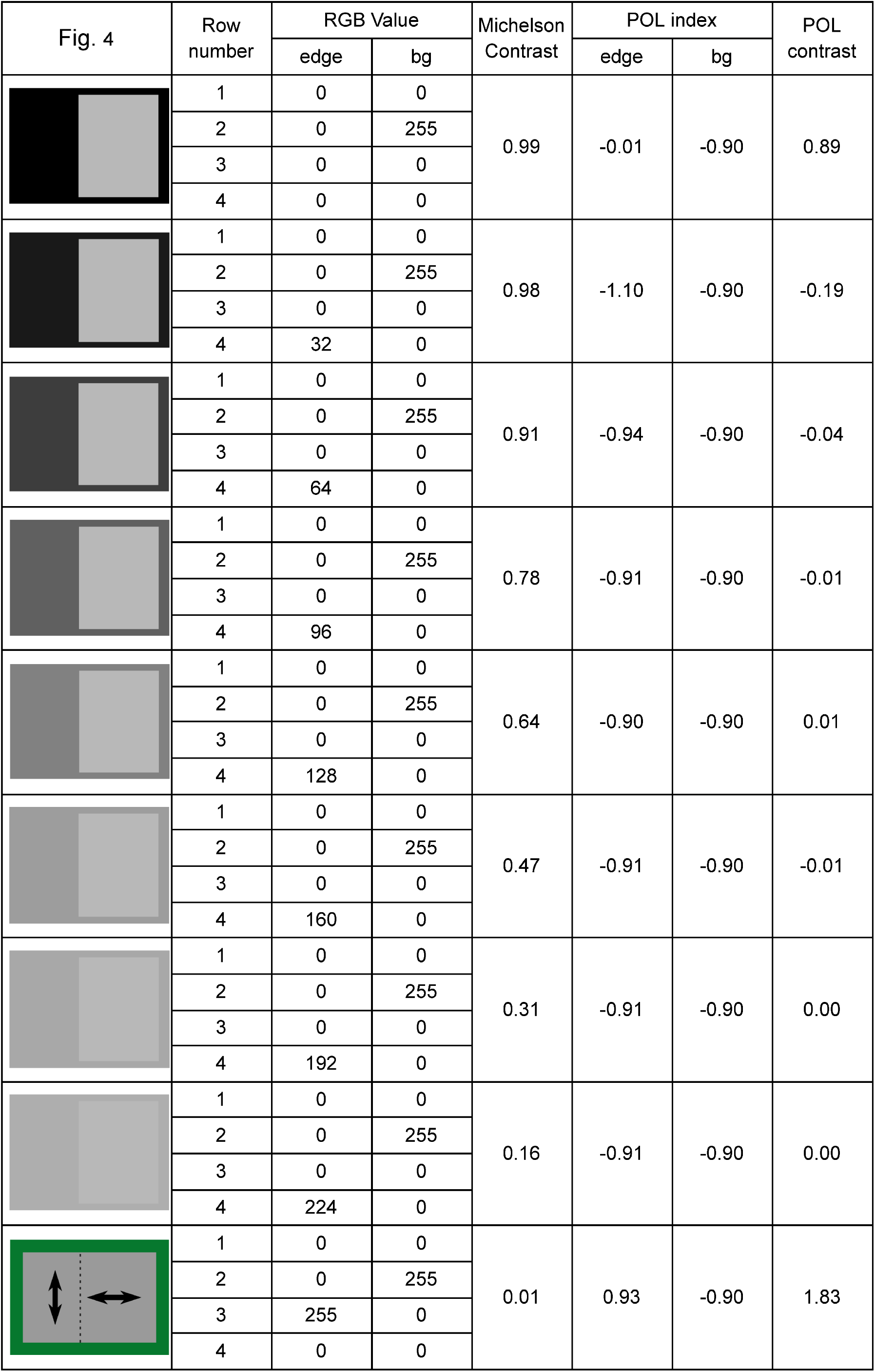
Stimuli characteristics for the experiment of Fig. 4. For each experimental condition, the RGB value of each row, the Michelson contrast between edge and background, the Polarization index (see Intensity and polarization screen), and the Polarization contrast (see Visual stimuli) are specified. The abbreviation bg stands for background.

| Fig. 4 | Row number | RGB Value |  | Michelson Contrast | POL index |  | POL contrast |
| --- | --- | --- | --- | --- | --- | --- | --- |
|  |  | edge | bg |  | edge | bg |  |
|  | 1 | 0 | 0 | 0.99 | -0.01 | -0.90 | 0.89 |
|  | 2 | 0 | 255 |  |  |  |  |
|  | 3 | 0 | 0 |  |  |  |  |
|  | 4 | 0 | 0 |  |  |  |  |
|  | 1 | 0 | 0 | 0.98 | -1.10 | -0.90 | -0.19 |
|  | 2 | 0 | 255 |  |  |  |  |
|  | 3 | 0 | 0 |  |  |  |  |
|  | 4 | 32 | 0 |  |  |  |  |
|  | 1 | 0 | 0 | 0.91 | -0.94 | -0.90 | -0.04 |
|  | 2 | 0 | 255 |  |  |  |  |
|  | 3 | 0 | 0 |  |  |  |  |
|  | 4 | 64 | 0 |  |  |  |  |
|  | 1 | 0 | 0 | 0.78 | -0.91 | -0.90 | -0.01 |
|  | 2 | 0 | 255 |  |  |  |  |
|  | 3 | 0 | 0 |  |  |  |  |
|  | 4 | 96 | 0 |  |  |  |  |
|  | 1 | 0 | 0 | 0.64 | -0.90 | -0.90 | 0.01 |
|  | 2 | 0 | 255 |  |  |  |  |
|  | 3 | 0 | 0 |  |  |  |  |
|  | 4 | 128 | 0 |  |  |  |  |
|  | 1 | 0 | 0 | 0.47 | -0.91 | -0.90 | -0.01 |
|  | 2 | 0 | 255 |  |  |  |  |
|  | 3 | 0 | 0 |  |  |  |  |
|  | 4 | 160 | 0 |  |  |  |  |
|  | 1 | 0 | 0 | 0.31 | -0.91 | -0.90 | 0.00 |
|  | 2 | 0 | 255 |  |  |  |  |
|  | 3 | 0 | 0 |  |  |  |  |
|  | 4 | 192 | 0 |  |  |  |  |
|  | 1 | 0 | 0 | 0.16 | -0.91 | -0.90 | 0.00 |
|  | 2 | 0 | 255 |  |  |  |  |
|  | 3 | 0 | 0 |  |  |  |  |
|  | 4 | 224 | 0 |  |  |  |  |
|  | 1 | 0 | 0 | 0.01 | 0.93 | -0.90 | 1.83 |
|  | 2 | 0 | 255 |  |  |  |  |
|  | 3 | 255 | 0 |  |  |  |  |
|  | 4 | 0 | 0 |  |  |  |  |

**Table S4.**
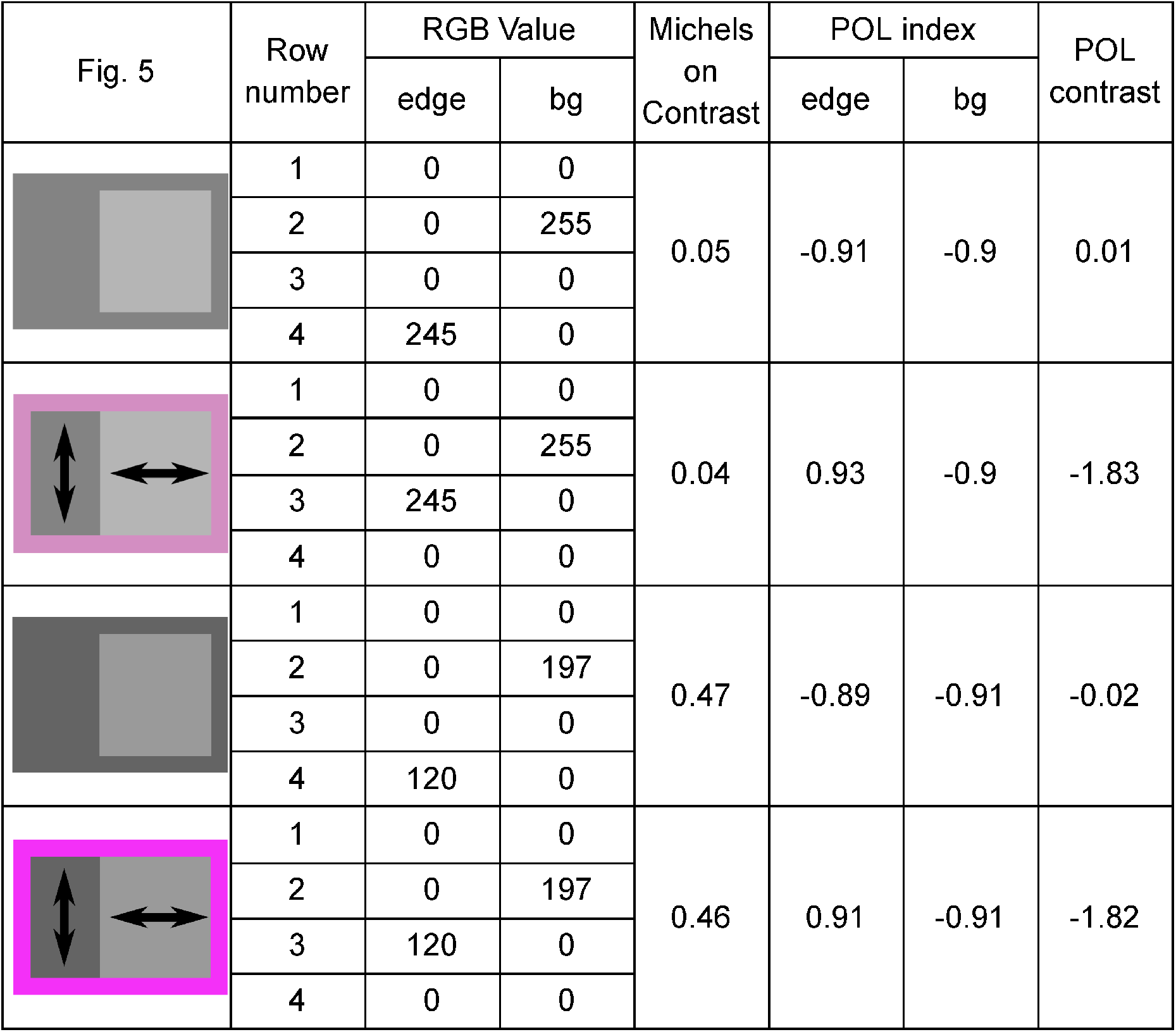
Stimuli characteristics for the experiment of Fig. 5. For each experimental condition, the RGB value of each row, the Michelson contrast between edge and background, the Polarization index (see Intensity and polarization screen), and the Polarization contrast (see Visual stimuli) are specified. The abbreviation bg stands for background.

| Fig. 5 | Row number | RGB Value |  | Michelson Contrast | POL index |  | POL contrast |
| --- | --- | --- | --- | --- | --- | --- | --- |
|  |  | edge | bg |  | edge | bg |  |
|  | 1 | 0 | 0 | 0.05 | -0.91 | -0.9 | 0.01 |
|  | 2 | 0 | 255 |  |  |  |  |
|  | 3 | 0 | 0 |  |  |  |  |
|  | 4 | 245 | 0 |  |  |  |  |
|  | 1 | 0 | 0 | 0.04 | 0.93 | -0.9 | -1.83 |
|  | 2 | 0 | 255 |  |  |  |  |
|  | 3 | 245 | 0 |  |  |  |  |
|  | 4 | 0 | 0 |  |  |  |  |
|  | 1 | 0 | 0 | 0.47 | -0.89 | -0.91 | -0.02 |
|  | 2 | 0 | 197 |  |  |  |  |
|  | 3 | 0 | 0 |  |  |  |  |
|  | 4 | 120 | 0 |  |  |  |  |
|  | 1 | 0 | 0 | 0.46 | 0.91 | -0.91 | -1.82 |
|  | 2 | 0 | 197 |  |  |  |  |
|  | 3 | 120 | 0 |  |  |  |  |
|  | 4 | 0 | 0 |  |  |  |  |

**Table S5.**
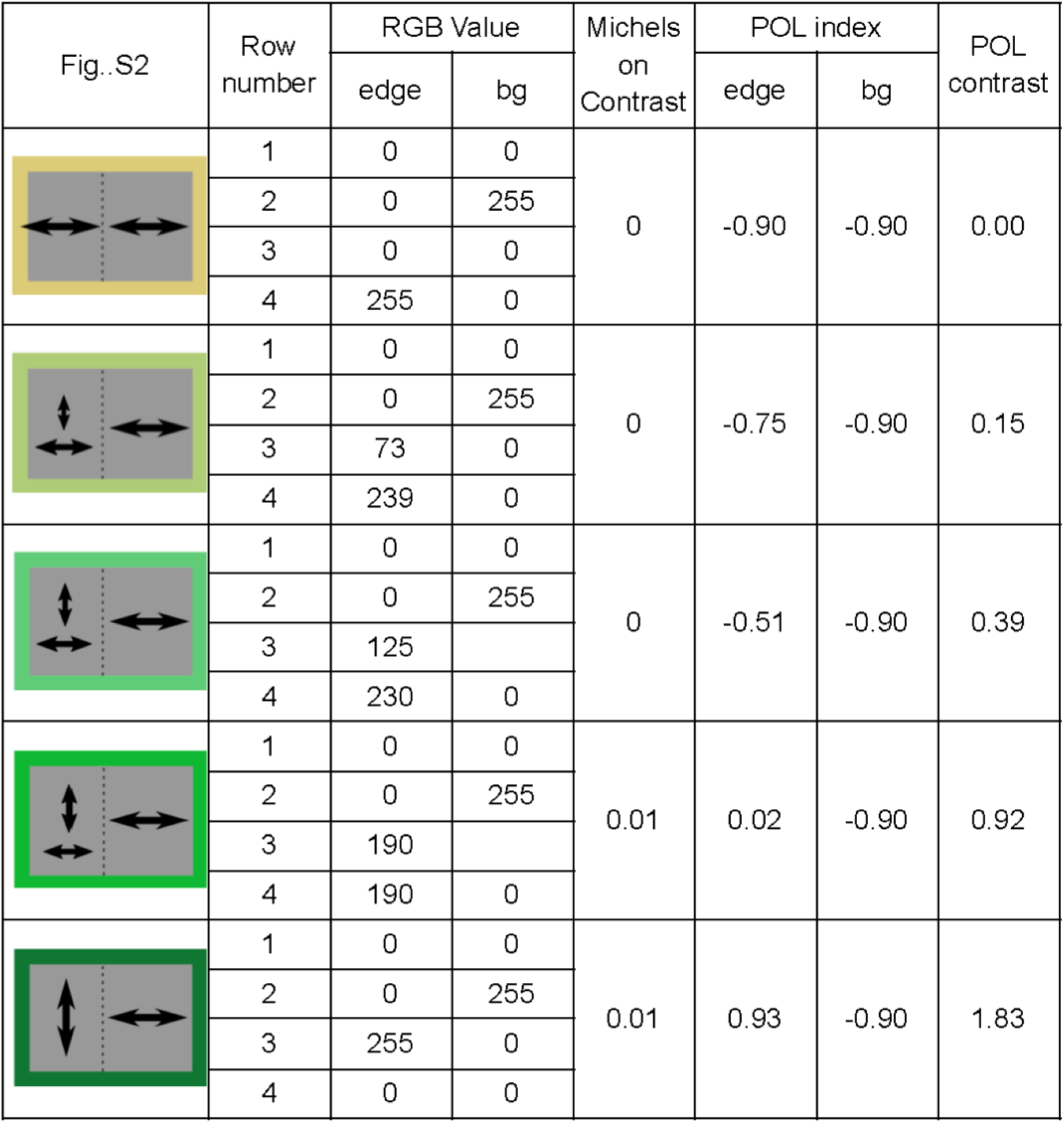
Stimuli characteristics for the experiment of Fig. S2. For each experimental condition, the RGB value of each row, the Michelson contrast between edge and background, the Polarization index (see Intensity and polarization screen), and the Polarization contrast (see Visual stimuli) are specified. The abbreviation bg stands for background.

| Fig..S2 | Row number | RGB Value |  | Michels on Contrast | POL index |  | POL contrast |
| --- | --- | --- | --- | --- | --- | --- | --- |
|  |  | edge | bg |  | edge | bg |  |
|  | 1 | 0 | 0 | 0 | -0.90 | -0.90 | 0.00 |
|  | 2 | 0 | 255 |  |  |  |  |
|  | 3 | 0 | 0 |  |  |  |  |
|  | 4 | 255 | 0 |  |  |  |  |
|  | 1 | 0 | 0 | 0 | -0.75 | -0.90 | 0.15 |
|  | 2 | 0 | 255 |  |  |  |  |
|  | 3 | 73 | 0 |  |  |  |  |
|  | 4 | 239 | 0 |  |  |  |  |
|  | 1 | 0 | 0 | 0 | -0.51 | -0.90 | 0.39 |
|  | 2 | 0 | 255 |  |  |  |  |
|  | 3 | 125 |  |  |  |  |  |
|  | 4 | 230 | 0 |  |  |  |  |
|  | 1 | 0 | 0 | 0.01 | 0.02 | -0.90 | 0.92 |
|  | 2 | 0 | 255 |  |  |  |  |
|  | 3 | 190 |  |  |  |  |  |
|  | 4 | 190 | 0 |  |  |  |  |
|  | 1 | 0 | 0 | 0.01 | 0.93 | -0.90 | 1.83 |
|  | 2 | 0 | 255 |  |  |  |  |
|  | 3 | 255 | 0 |  |  |  |  |
|  | 4 | 0 | 0 |  |  |  |  |

## Notes

### Competing Interest Statement

The authors have declared no competing interest.

